# CNV Neurons Are Rare in Aged Human Neocortex

**DOI:** 10.1101/303404

**Authors:** William D. Chronister, Margaret B. Wierman, Ian E. Burbulis, Matthew J. Wolpert, Mark F. Haakenson, Joel E. Kleinman, Thomas Hyde, Daniel R. Weinberger, Stefan Bekiranov, Michael J. McConnell

**Affiliations:** Departments of Biochemistry and Molecular Genetics, University of Virginia School of Medicine, Charlottesville, VA 22908; Departments of Neuroscience, University of Virginia School of Medicine, Charlottesville, VA 22908; Centers for Public Health Genomics, University of Virginia School of Medicine, Charlottesville, VA 22908; Brain Immunology and Glia, University of Virginia School of Medicine, Charlottesville, VA 22908; Children’s Health Research, University of Virginia School of Medicine, Charlottesville, VA 22908; Lieber Institute for Brain Development, Johns Hopkins Medical Campus, Baltimore, MD 21205; Department of Psychiatry, Johns Hopkins University School of Medicine, Baltimore, MD; Department of Neurology, Johns Hopkins University School of Medicine, Baltimore, MD; Departments of Neuroscience and the McKusick-Nathans Institute of Genetic Medicine, Baltimore, MD.

## Abstract

Megabase-scale somatic copy number variants (CNVs) alter allelic diversity in a subset of human neocortical neurons. Reported frequencies of CNV neurons range from ∼5% of neurons in some individuals to greater than 30% in other individuals. Genome-wide and familial studies implicitly assume a constant brain genome when assessing the genetic risk architecture of neurological disease, thus it is critical to determine whether divergent reports of CNV neuron frequency reflect normal individual variation or technical differences between approaches. We generated a new dataset of over 800 human neurons from 5 neurotypical individuals and developed a computational approach that measures single cell library quality based on Bayesian Information Criterion and identifies integer-like variant segments from population-level statistics. A brain CNV atlas was assembled using our new dataset and published data from 10 additional neurotypical individuals. This atlas reveals that the frequency of neocortical CNV neurons varies widely among individuals, but that this variability is not readily accounted for by tissue quality or CNV detection approach. Rather, the age of the individual is anti-correlated with CNV neuron frequency. Fewer CNV neurons are observed in aged individuals than young individuals.

## Introduction

Neocortical neurons are among the most diverse and longest-lived mammalian cells. The mammalian cerebral cortex is often put forward as a pinnacle of evolutionary complexity, and human-specific brain phenotypes are attributed to neocortical expansion during evolution (Geschwind and Rakic, 2013; Lui et al., 2011). Aberrant development of neocortical circuits is likewise associated with neuropsychiatric and neurodegenerative disease (Del Pino et al., 2018; Sudhof, 2017). Neurostereological approaches count ∼ 20 million neurons and ∼ 35 million glia in the human cerebral cortex (Pakkenberg et al., 2003) and single cell transcriptomic approaches are beginning to comprehensively catalogue human cortical diversity (Lake et al., 2016; Nowakowski et al., 2017). After decades of debate, it is now clear that human neocortical neurons are not normally regenerated during human lifespan (Bhardwaj et al., 2006; Rakic, 2006). With some exceptions (Spitzer, 2017), neuronal cell types are also generally thought to be stable throughout life, but neuronal genomes are, perhaps, surprisingly labile.

Genome maintenance is critical in neurons, and enhanced DNA repair is a pharmacological target to ameliorate cognitive decline (Chow and Herrup, 2015; Madabhushi et al., 2014; McKinnon, 2017). However, somatic genomes are not stable (Frank, 2010; Lynch, 2010), and somatic mutations are common in neurotypical human brains (McConnell et al., 2017). Brain somatic mosaicism has been most widely assessed in prefrontal cerebral cortex, where the impact of a small population of aberrant neurons can be significant, and cross-sectional studies of post-mortem neurotypical individuals are not confounded by cell turnover. Collectively, these studies suggest that every neocortical neuron may contain private somatic variants. Single nucleotide variants (SNVs) are especially common with hundreds per neuron reported (Bae et al., 2017; Lodato et al., 2015) and with frequencies of over 3000 SNVs per neuron observed in aged individuals (Lodato et al., 2017). Endogenous mobile elements such as LINE1 retrotransposons are also active during brain development (Coufal et al., 2009; Muotri et al., 2005). Reported frequencies of *de novo* mobile element events range from <1 to >7 per neuron (Baillie et al., 2011; Evrony et al., 2012; Upton et al., 2015). Mobile element activity has also been linked to the generation of copy number variants (CNVs) (Erwin et al., 2016; Gilbert et al., 2002). Large mosaic CNVs duplicate or delete several megabases (Mb) of genomic sequence in a subpopulation of neocortical neurons (Cai et al., 2014; Knouse et al., 2016; McConnell et al., 2013; Piotrowski et al., 2008). By contrast to other brain somatic variants which rarely impact protein-coding sequence, gene density in the human genome is >10 genes per Mb. Reanalysis of published data (Cai et al., 2014; Knouse et al., 2014; McConnell et al., 2013; van den Bos et al., 2016) herein found an average of 63 genes affected per neuronal CNV.

During the past decade, large CNVs have been recognized as major contributors to human genetic diversity (Conrad et al., 2010; Lupski, 2015; Redon et al., 2006). At the population level, SNVs are collectively more numerous than CNVs, but CNVs impact an order of magnitude more genome sequence (∼10%), and some CNVs show evidence of positive selection during human evolution (Perry et al., 2007; Sudmant et al., 2015; Zarrei et al., 2015). In individuals, *de novo* CNVs represent rare variants with a strong contribution to the genetic risk of schizophrenia, autism, and other neurological disorders (Fromer et al., 2014; Iossifov et al., 2014; Marshall et al., 2017; Morrow, 2010; Sebat et al., 2007). Whereas the consequences of germline CNVs have been inferred from population level studies, neuronal CNV studies to date have been underpowered to determine if the genes affected by neuronal CNVs contribute to brain development, function, and disease.

We sought to determine how neuronal CNVs shape the genetic architecture of the cerebral cortex by assembling an atlas of brain somatic CNVs from 15 neurotypical individuals. A new dataset of 827 human cerebral cortical nuclei from 5 neurotypical individuals was generated using a single whole genome amplification (WGA) approach. Read depth-based single cell CNV detection is validated by studies that observe clonal evolution in tumor populations and reciprocal genomic events among sister cells in 8-cell human embryos (Navin et al., 2011; Vanneste et al., 2009). However, large neuronal CNVs are rarely clonal and virtually impossible to confirm in bulk tissue; thus, we also developed an unbiased CNV detection approach based on population-level statistics and established the neuronal CNV atlas with 295 neuronal CNVs. Published datasets (Cai et al., 2014; Knouse et al., 2014; McConnell et al., 2013; van den Bos et al., 2016) from 10 other neurotypical individuals added 212 neuronal CNVs to the atlas. Initial analysis of the neuronal CNV atlas finds substantial interindividual variability in the frequency of CNV neurons, but also supports the hypothesis that some long genomic loci shape the genetic architecture of neurotypical human brains.

## Results

### Generation of Single Cell Neuronal and Non-neuronal Genomic Data

We obtained prefrontal cortex from five male individuals aged 0.36, 26, 49, 86, and 95 years (Table 1) and performed fluorescence-activated nuclei sorting (FANS) to isolate single neuronal and non-neuronal nuclei (Fig. 1A). WGA was performed using PicoPLEX (Rubicon Genomics), an approach similar to multiple annealing and loop-based amplification (MALBAC) (Zong et al., 2012) that produces Illumina-compatible libraries with 48 unique barcodes. Prior to library pooling and sequencing on Illumina-based platforms, we found that ∼60% of WGA reactions produced measurable product. Single-end or paired-end sequencing (50, 75, or 100bp) of 48 pooled libraries on Illumina HiSeq Rapid platforms or of smaller pools (<17 libraries) on MiSeq platforms routinely produced more than 1 million reads per library after duplicate removal. Neither paired-end nor longer read sequencing altered data quality. Reads were aligned to hg19 and read depth was calculated across 4,505 genomic bins, each containing ∼500kb of uniquely mappable sequence (mean bin size = 687kb +/− 1,072kb SD). Collectively, this approach (Fig. 1B) generated >134.7 Gb of genomic sequence from 829 single cell genome libraries (162.5 +/− 115.1 Mb/cell).

**Figure 1.**
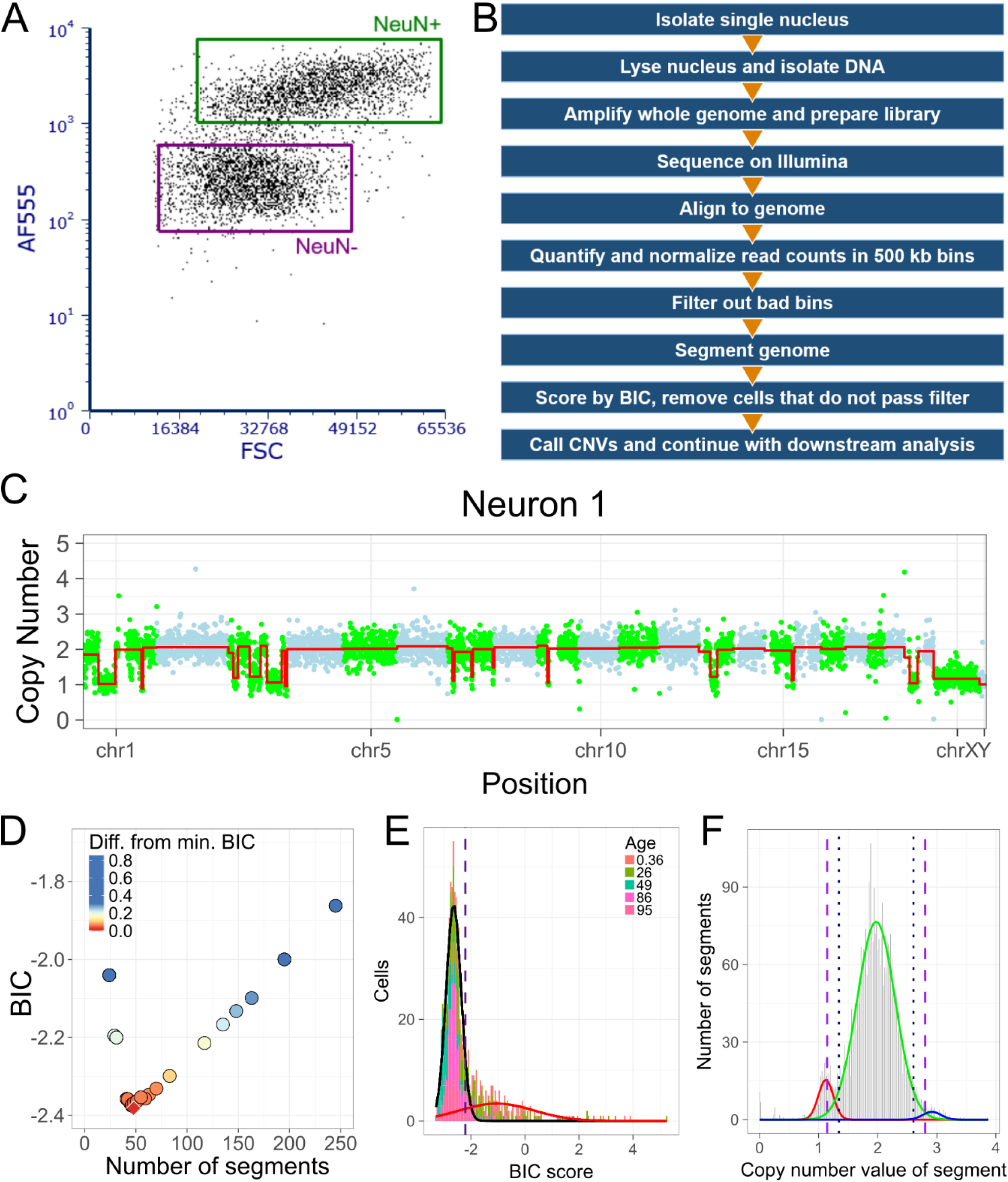
Optimization of sample quality filtering and CNV calling in a new single cell dataset. (**A**) Representative FANS plot of NeuN Immunostaining of brain nuclei (AF555). (**B**) Summary of single nuclei analysis pipeline. (**C**) CNV profile of test data Neuron 1. Genome is arranged horizontally by chromosome. Dots represent read depth-derived CN values of genomic bins and are colored by chromosome, alternating between green and light blue. Red line represents segmentation output from DNAcopy. (**D**) Comparison of BIC scores for segmentation of Neuron 1 using different settings of alpha, undoSD, and min.width. Red diamond represents lowest BIC score. (**E**) Histogra of BIC scores for entire single cell dataset. Gaussian distributions fit to data are depicted in black and red. Dashed line indicates BIC cutoff of < −2.21. (**F**) Histogram of segments 5–45 bins in size. Gaussian distributions fit to data are shown in red, green, and blue. Long dashed lines indicate stringent CNV cutoffs of < 1.14 and > 2.80. Short dashed lines indicate lenient CNV cutoffs of < 1.34 and > 2.60.

**Table 1.** Overview of PicoPLEX data.

| Age | Sex | Cell type | BIC pass/total cells (%) | Cells with CNV(s) (%), lenient | Cells with CNV(s) (%), stringent | Total CNVs (lenient) | Total CNVs (stringent) |
| --- | --- | --- | --- | --- | --- | --- | --- |
| 0.36 | Male | Neuron | 46/138 (33.3) | 14 (30.4) | 7 (15.2) | 41 | 12 |
| 26* | Male | Neuron | 108/184 (58.7) | 25 (23.1) | 15 (13.9) | 76 | 33 |
| 26* | Male | Non-neuron | 43/63 (68.3) | 2 (4.7) | 0 (0) | 2 | 0 |
| 49 | Male | Neuron | 99/101 (98.0) | 11 (11.1) | 8 (8.1) | 113 | 75 |
| 49 | Male | Non-neuron | 26/28 (92.9) | 2 (7.7) | 2 (7.7) | 4 | 2 |
| 86 | Male | Neuron | 101/118 (85.6) | 4 (4.0) | 3 (3.0) | 20 | 15 |
| 86 | Male | Non-neuron | 46/55 (83.6) | 4 (8.7) | 2 (4.3) | 9 | 3 |
| 95 | Male | Neuron | 120/140 (85.7) | 8 (6.7) | 5 (4.2) | 45 | 28 |

\*Same individual as 26 year old listed in Table 2.

^*^Same individual as 26 year old listed in Table 2.

### Optimization of Read Depth-based Single Cell Genomic Segmentation

Neuronal CNV detection is inherently challenging because one cannot know the state of the genome prior to WGA. Clonal neuronal CNVs are rare which generally precludes targeted sequencing approaches for validation in the source material. For this reason, CNV calling approaches have been employed conservatively with bias toward avoiding Type I error at the risk of Type II error. To optimize CNV detection, we selected a test dataset of six representative single cell WGA libraries from five neurons and one trisomy fibroblast with varied subjective quality (Fig. 1C, Fig. S1A-E) and assessed DNAcopy (Olshen et al., 2004) - a CNV segmentation tool - performance across algorithm parameter space by calculating Bayesian Information Criterion (BIC)(Schwarz, 1978). Given that DNAcopy was shown to be susceptible to calling false positive CNVs containing strong autocorrelated noise (Muggeo and Adelfio, 2011), we simulated model cells that contained either no CNVs but strong autocorrelated noise (NULL model) or known CNVs with weak residual autocorrelated noise (ALT model) based on the read-depth statistics of the original cell (Fig. S1G). Finally, we applied optimal parameters to all 829 libraries, rejected 238 due to poor BIC scores, and assessed specificity and sensitivity based on monosomy X detection in 589 male neural nuclei.

**Figure S1.**
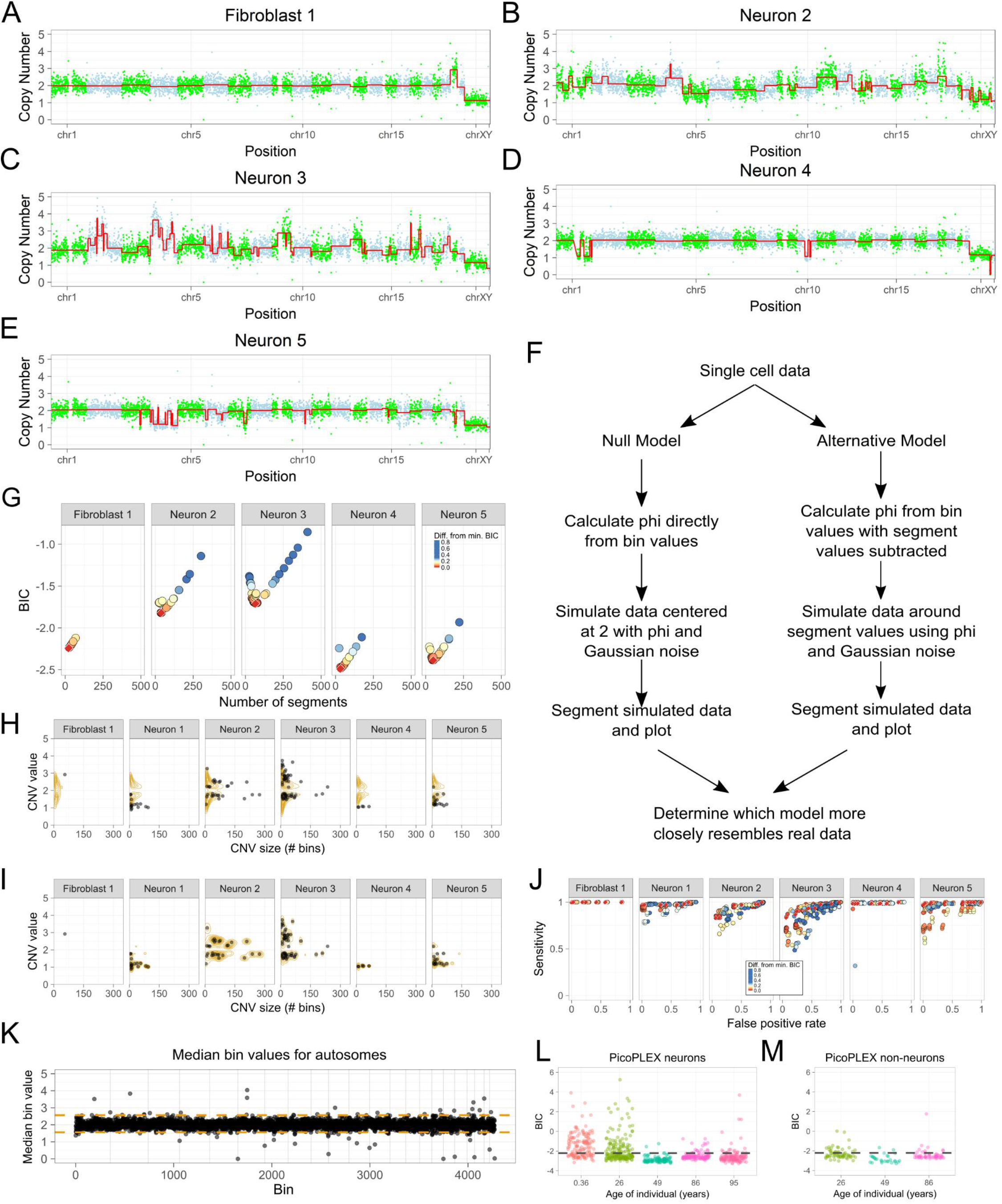
(related to Figure 1). Optimization of sample filtration and segmentation. (**A-E**) CNV profile of test data Fibroblast 1 (**A**) and Neurons 2–5 (B-E). Genome is arranged horizontally by chromosome. Read depth-derived CN values of genomic bins and are colored by chromosome, alternating between green and light blue. Red line represents segmentation output from DNAcopy. (**F**) Flowchart summarizing simulation of data based on authentic single cell data using the null and alternative models. (**G**) Comparison of BIC scores for segmentation of Fibroblast 1 and Neurons 2–5 using different settings of alpha, undoSD, and min.width. A red diamond is used to highlight the lowest BIC score for each cell. (**H**) Comparison of CNVs detected in real data (black dots) to CNVs detected in simulated data (yellow contours) under the null and (**I**) alternative models across six test cells. For the purposes of these tests, thresholds of CN < 1.825 for deletion and CN > 2.175 for duplication were used. (**J**) ROC curves demonstrating the sensitivity and false positive rates of CNV detection under different DNAcopy settings. Points are colored according to difference from minimum BIC score for each cell, as in panel G; similarly, red diamonds are used to highlight the best BIC scores for each cell. 10 CNV thresholds were used, ranging from < 1.825 for deletion and > 2.175 for duplication to < 1.99 for deletion and > 2.01 for duplication. For that reason, BIC scores from the same segmentation may appear up to 10 times in each plot depending on changes in sensitivity or false positive rate resulting from changes in the CNV thresholds used. (**K**) Median bin values for autosomal bins in PicoPLEX data. Orange dashed lines represent bin exclusion thresholds of > 2.57 and < 1.56, as determined by Tukey’s outlier test. (**L**) BIC scores for PicoPLEX neurons and (**M**) non-neurons. Dashed line indicates BIC cutoff of < −2.21.

BIC scores provide a log-likelihood estimate of the segmentation algorithm’s performance by penalizing both the variance within a segment and the number of called segments. Thus, low BIC scores indicate the best algorithm performance on any given single cell library. Parameter space in DNAcopy is defined by three user-tunable parameters: significance threshold (alpha), minimum number of genomic bins required to call a copy number state (min.width) and the number of standard deviations between the levels of copy number states in order to maintain the copy number state change (undoSD). Using the test dataset, we calculated BIC scores for dozens of combinations of these parameters and found that settings of alpha=0.001, min.width=5, and undoSD=0 consistently produce the lowest or near-lowest BIC scores for each library (Fig. 1D and S1G). These parameters identified monosomy X in all 6 male cells, and trisomy 21 in only the fibroblast. We also observed that minimum BIC scores were lower in WGA libraries with less overall bin-to-bin variation in read-depth (Fig. 1C, S1A-E vs. Fig. S1G), suggesting that low BIC scores are associated with high quality WGA reactions.

We assessed the extent to which segmentation algorithm performance was impaired by Gaussian or autocorrelated noise using 200 simulated datasets from each of the 6 test cells (Fig. S1F). Simulated single cell datasets were built based on fitting two models’ parameters to observed read-depth data (e.g., the standard deviation and a measure of correlated noise strength, φ). Estimation of φ is essential given that autocorrelated noise can be a source of false positive calls by DNAcopy (Muggeo and Adelfio, 2011). NULL model cells contained no “true” CNVs and were generated from bin-level test data using an autoregressive model of order 1 (i.e., AR(1)) whereby the *n^th^* bin value depends on the *n−1^th^* bin value with a coupling strength 9 plus Gaussian noise. ALT model cells contained only “true” CNVs based on DNAcopy output plus residual correlated and Gaussian noise. The synthetic NULL and ALT cells were assessed for CNV event sizes and CN state relative to the real data (Fig. S1 H-I). CNV sizes and values derived from segmentation of the ALT model cells, but not the NULL model cells, match the experimental data well (Fig. S1H, I). Thus, we conclude that the single cell neuronal genomic data are more consistent with actual CNV events plus Gaussian noise (i.e., ALT model) than autocorrelated and Gaussian noise (i.e., NULL model). This suggests that the DNAcopy parameters identified can produce low false positives in single cell genomic data. Consistent with this, the DNAcopy parameters which yielded the lowest BIC scores, also produced the highest sensitivity and specificity based on concordance with ALT model cells (Fig. S1J).

When segmentation was performed on the entire dataset, we observed that some genomic bins routinely deviated from median read depth, and that these bins interfered with segmentation and CNV identification (Fig. S1K). To objectively exclude aberrant bins, we performed Tukey’s outlier test on the median copy number (CN) values of all 4,505 bins across 829 genomic datasets and removed 101 (2.2%) outlier bins from subsequent analysis. Given that aberrant bins directly affect BIC scores, BIC scores presented previously were calculated after outlier bin removal. To objectively exclude WGA libraries that would be most prone to poor algorithm performance, we fit a two-Gaussian mixture model to the BIC score data and used the 95^th^ percentile of the low BIC score Gaussian distribution to arrive at a BIC score < −2.21 as a cutoff for high quality samples (Fig. 1E). This resulted in 589/829 cells passing the BIC cutoff (71.3%), including 474/681 (69.6%) NeuN+ nuclei and 115/146 (78.8%) NeuN- nuclei (Fig. S1 L-M).

The veracity of CNV identification based on read depth has previously been assessed by adherence to expected near-integer CN states (Garvin et al., 2015; Knouse et al., 2016; McConnell et al., 2013); however, the data display and DNAcopy assigns non-integer CN values to detected segments. To objectively identify integer-like segments in the dataset, we fit the distribution of CN values to a Gaussian mixture model containing three Gaussians (Fig. 1F). Consistent with integer-like expectations of true positive CNV calls, we found that the distributions displayed two modes occurring near CN values of 2 (euploid) and 1 (deletion of a chromosomal segment). In addition, we observed a heavy tail near CN values around 3 (duplication of a chromosomal segment). The fitted Gaussian means were 1.12, 1.97 and 2.92 (see Methods for details), which are consistent with expected monosomy, disomy, and trisomy values. We then used the middle Gaussian with mean 1.97 (i.e., near the CN = 2 state) as a NULL model of no change and defined lenient CN loss and gain thresholds based on a two-tailed p-value ≤ 0.05 (≤ 1.34 for a deletion and ≥ 2.60 for a duplication) and stringent loss and gain thresholds based on a two-tailed p-value ≤ 0.01 (≤ 1.14 for a deletion and ≥ 2.80 for a duplication). Using monosomy X in 589 male nuclei as a positive control, we found that the stringent threshold led to more false negatives (47.2%) than the lenient threshold (0.5%). Importantly, we cannot exclude the possibility that some false negatives are due to true X chromosome CNVs or disomy X in male genomes. We present results using the lenient 0.05 p-value thresholds and that of the stringent in supplementary figures. Notably, the two sets of results are broadly concordant.

### Contribution of Mosaic CNVs to Neuronal Diversity

Large CNVs inevitably alter the CN state of a brain-expressed gene, which is not surprising given that more of the genome is expressed in neurons than in other cell types (Uhlen et al., 2016). Our analysis (Table 1) identified 310 CNVs in 70 of 589 neural genomes (11.9%). CNVs ranged in size from 2.9Mb to 159.1Mb with a mean of 16.5 Mb and standard deviation of 20.1 Mb (Fig. 2A). With stringent criteria that are prone to false negatives, we still identify 168 CNVs (mean 18.0 +/− 22.3 MB) in 42 neural nuclei (7.1%) (Fig. S2A). CNVs were detected in each individual examined and, given their size and frequency (Fig. 2B, Fig. S2D), represent a clear contribution to the genetic architecture of the brain (Fig. 2C, Fig. S2G). We note that much smaller percentages of phenotypically distinct cells can bring about focal epilepsies (Marin-Valencia et al., 2014) and are essential for normal brain function (e.g., adult-born dentate granule neurons (Anacker and Hen, 2017; Christian et al., 2014)).

**Figure 2.**
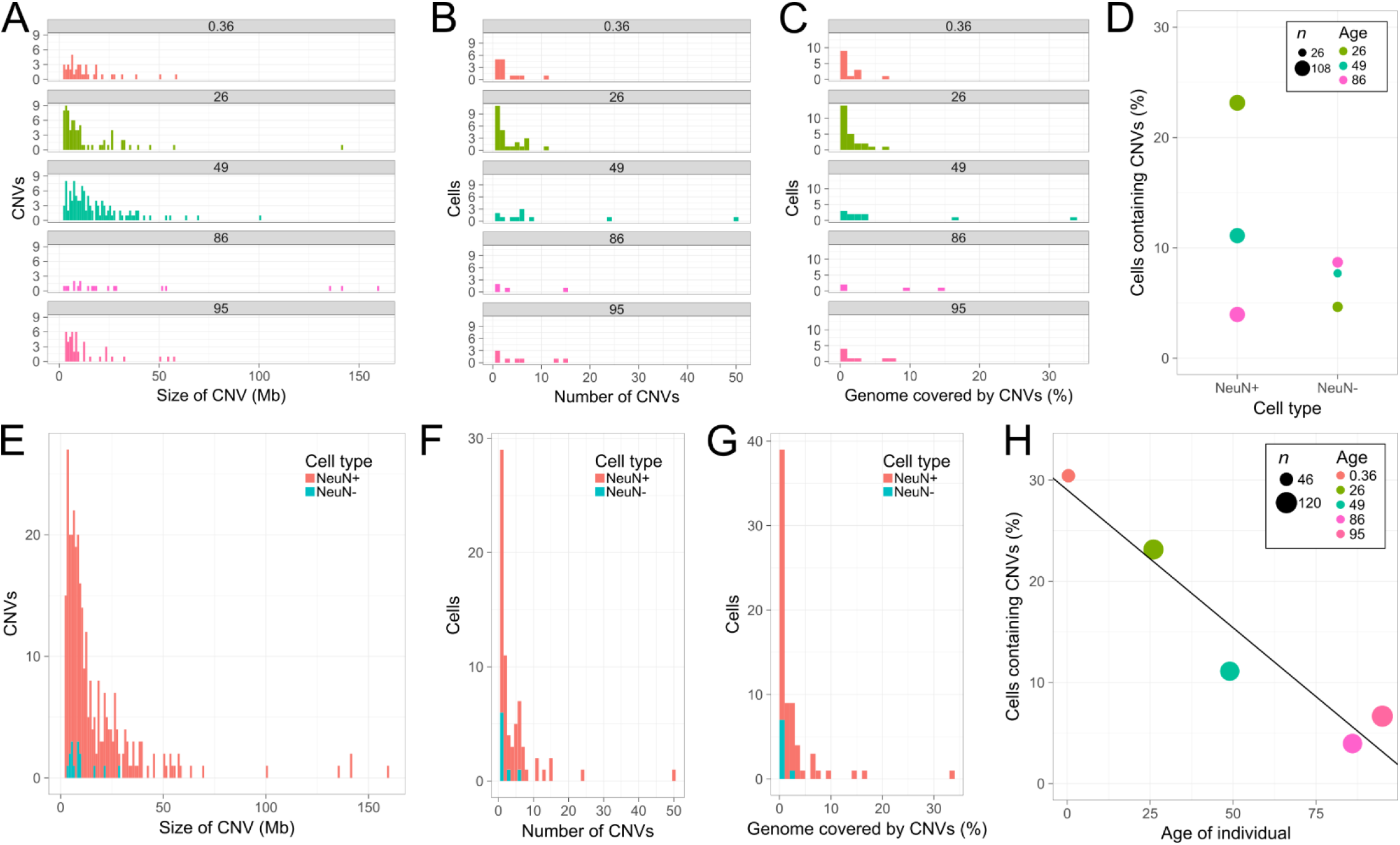
Contribution of mosaic CNVs to neuronal diversity across individuals. (**A**) Histogram showing CNV sizes among the neurons from five individuals aged 0.36 to 95. (**B**) Histogram showing number of CNVs in CNV neurons from five individuals. Neurons with 0 CNVs were excluded from this plot. (**C**) Histogram showing percent genome coverage by CNVs in CNV neurons from five individuals. Neurons with 0 CNVs were excluded from this plot. (**D**) Comparison of percent cells containing CNVs between NeuN+ and NeuN- cells from three individuals. (**E**) Comparison of CNV size between NeuN+ and NeuN- cells. (**F**) Comparison of number of CNVs between NeuN+ and NeuN- cells. Cells with 0 CNVs were excluded from this plot. (**G**) Comparison of percent genome coverage by CNVs between NeuN+ and NeuN- cells. Cells with 0 CNVs were excluded from this plot. (**H**) Percent of neurons containing CNVs plotted against the age of the individual from which they were collected (linear fit, multiple R^2^ = 0.9224, p < 0.01). Lenient CNV detection thresholds were used for all of the above panels.

**Figure S2.**
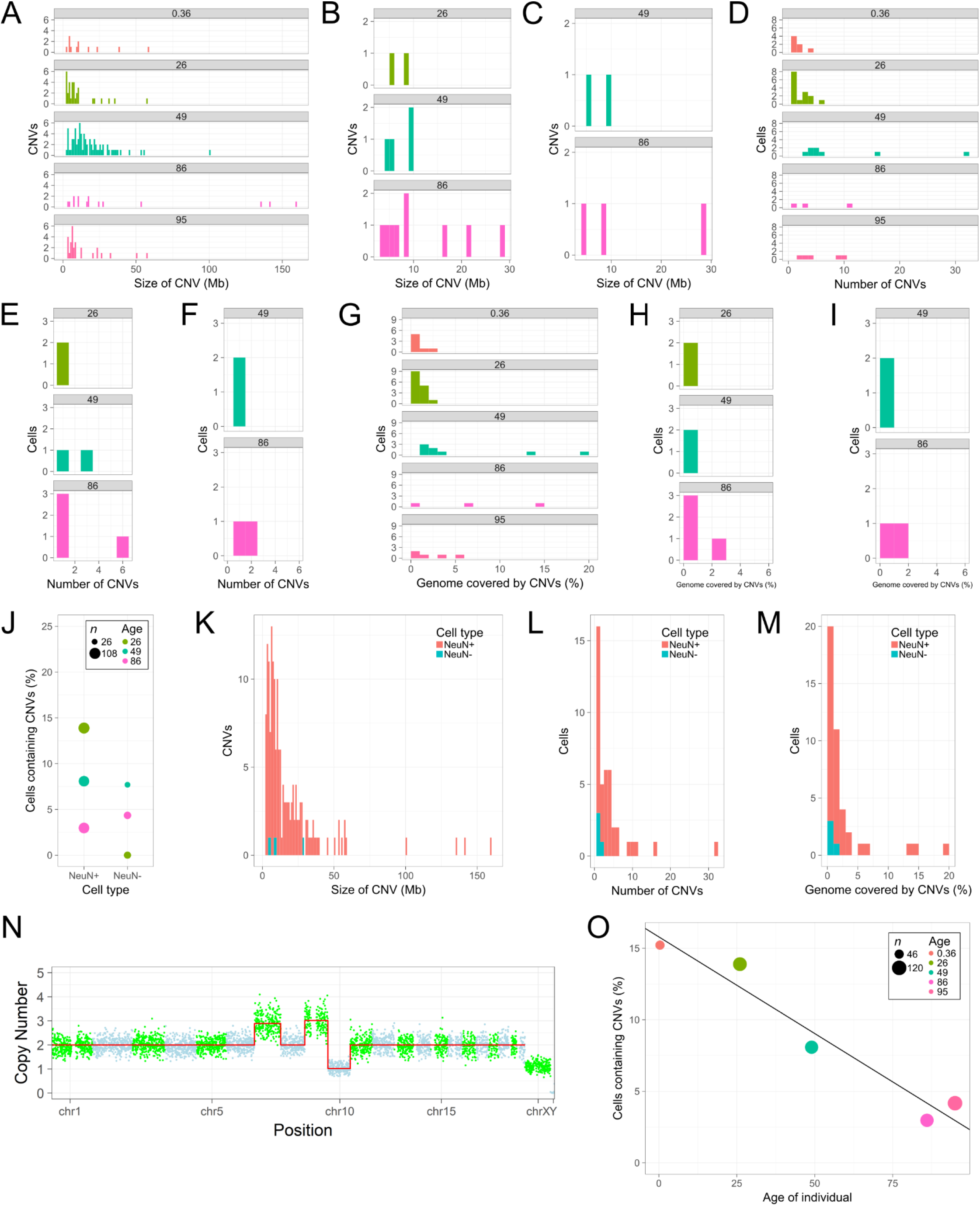
(related to Figure 2). Contribution of mosaic CNVs to brain cell diversity across individuals and CNV thresholds. (**A**) Histogram showing CNV sizes among the neurons from five individuals aged 0.36 to 95. Stringent thresholds were used. (**B and C**) Histograms showing CNV sizes among the non-neurons from three individuals aged 49 to 86. CNVs were detected using lenient thresholds (**B**) and stringent thresholds (**C**). (**D**) Histogram showing number of CNVs in CNV neurons from five individuals. Neurons with 0 CNVs were excluded from this plot. Stringent thresholds. (**E and F**) Histograms showing number of CNVs in CNV non-neurons from three individuals. Non-neurons with 0 CNVs were excluded from this plot. CNVs were detected using lenient thresholds (**E**) and stringent thresholds (F). (**G**) Histogram showing percent genome coverage by CNVs in CNV neurons from five individuals. Neurons with 0 CNVs were excluded from this plot. Stringent thresholds. (H and I) Histogram showing percent genome coverage by CNVs in CNV non-neurons from three individuals. Non-neurons with 0 CNVs were excluded from this plot. CNVs were detected using lenient thresholds (**H**) and stringent thresholds (I). (**J**) Comparison of percent cells containing CNVs between NeuN+ and NeuN- cells from three individuals. Stringent thresholds. (**K**) Comparison of CNV size between NeuN+ and NeuN- cells. Stringent thresholds. (**L**) Comparison of number of CNVs between NeuN+ and NeuN- cells. Cells with 0 CNVs were excluded from this plot. Stringent thresholds. (**M**) Comparison of percent genome coverage by CNVs between NeuN+ and NeuN- cells. Cells with 0 CNVs were excluded from this plot. Stringent thresholds. (**N**) Neuron from 86 year old male displaying trisomy 7, trisomy 9, and monosomy 10. The cell also appears to show a Y chromosome loss. Red lines represent the Cn states at each bin across the genome; CNV calls are represented at the precise Cn determined by DNAcopy while euploid regions are depicted at exactly CN 2. (**O**) Percent of neurons containing CNVs plotted against the age of the individual from which they were collected (linear fit, multiple R^2^ = 0.9434, p < 0.01). Stringent thresholds.

We find that both neuronal and non-neuronal genomes harbor Mb-scale CNVs, consistent with previous reports (Cai et al., 2014; Knouse et al., 2016). Neurons harboring large CNVs (CNV neurons) were observed at different frequencies in each of three individuals from which non-neuronal cells were also analyzed (Fig. 2D). In the 26 year-old, 23.1% of neurons (25/108) were CNV neurons whereas only 4.7% of non-neuronal cells (2/43) contained CNVs. Fewer CNV neurons were observed in the 49 year-old (11.1%) and the 86 year-old (4.0%), but non-neuronal CNVs were observed at similar frequencies in all three individuals (4.7–8.7%). Interestingly, neuronal CNVs tended to be larger (Fig. 2E), occur in greater numbers (Fig. 2F), and affect more of the genome than non-neuronal CNVs (Fig. 2G). We found that mean genome coverage of CNVs ranged from 1.3 to 6.0% in neurons but only from 0.2 to 0.8% in non-neurons. CNV neurons in the 49 year old individual had an average of 6.0% of their genome at non-euploid CN states (186 MB +/− 317 MB). When restricted to stringent thresholds, similar ratios of neuronal and non-neuronal CNVs were still observed, but mean differences in non-euploid CN states were amplified (Fig. S2K-M).

The 474 neuronal nuclei we analyzed represent the largest CNV dataset of neurotypical human brains generated by one laboratory using a single WGA approach to date. We identified 62 CNV neurons (13.1%) with a mean CNV size of 16.8 Mb. The individual with the highest mean CNV size (37.2 Mb) was 86 years old (Fig. 2A); one neuron contained two trisomies and one monosomy (Fig. S2N). An average of 4.8 CNVs per CNV neuron was observed, but this also varied among individuals. The 49 year old individual had the highest (10.3 CNVs/CNV neuron) CNV burden in CNV neurons (Fig. 2B, C). A statistically significant linear decline in abundance of CNV neurons was observed with age (R^2^ = 0.9224; p = 0.0094) (Fig. 2H). The correlation was also significant at the stringent threshold (R^2^ = 0.9434; p = 0.0058) (Fig. S2O). These results suggest an increased CNV burden and a concordant decrease in the frequency of CNV neurons in aged human neocortex.

### Analysis and Integration of Publicly Available Data

Our data show that neurotypical human frontal cortex can contain as few as 4.0% CNV neurons in some individuals, but as many as 30.4% CNV neurons in other individuals. Four previous studies employing different WGA and analysis approaches have assessed additional individuals (Cai et al., 2014; Knouse et al., 2014; McConnell et al., 2013; van den Bos et al., 2016). In total, these datasets comprise 458 single neurons from 5 male and 5 female neurotypical individuals of different ages. The use of lab-specific WGA approaches and neuronal isolation techniques, as well as different CNV detection approaches, confounds direct comparison of new data with published results. Therefore, we obtained FASTQ files for publicly available data, harmonized these with our PicoPLEX dataset (e.g., alignment to hg19), assessed and excluded outlier bins (Fig. S3A-B) and evaluated WGA quality by calculating BIC scores (Fig. 3A-B, S3C-D) for each WGA approach.

**Figure 3.**
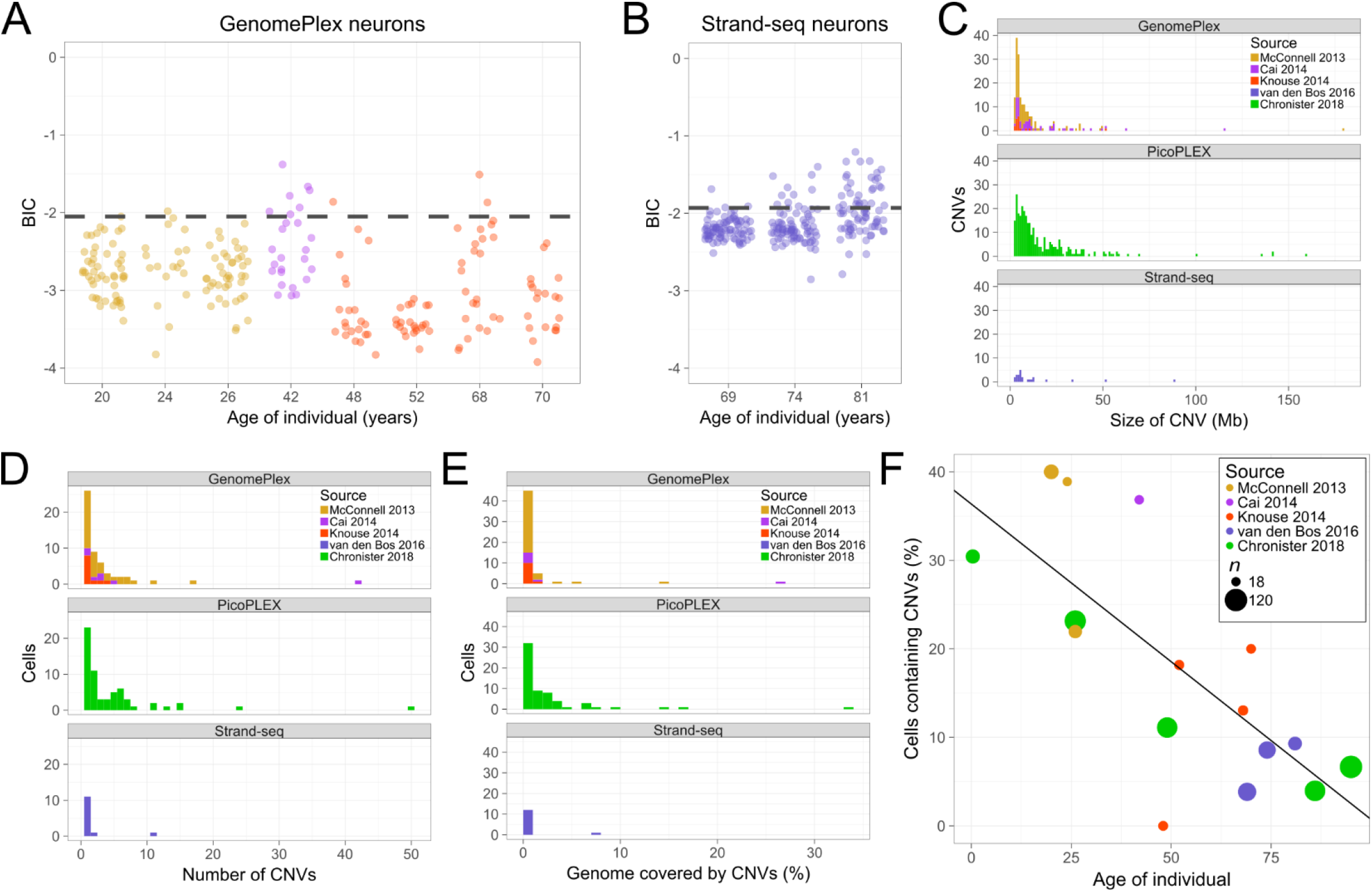
Integration of publicly available data supports trends seen in PicoPLEX data. (**A and B**) BIC scores for publicly available single neuron data amplified with GenomePlex (**A**) and Strand-seq (B). Black dashed lines indicate BIC cutoffs of < −2.05 (GenomePlex) and < −1.93 (Strand-seq). (**C**) Histogram showing CNV sizes among neurons across each WGA method and data source. (**D**) Histogram showing number of CNVs in each CNV neuron across each WGA method and data source. Neurons containing 0 CNVs were excluded from this plot. (**E**) Histogram showing percent genome coverage by CNVs in each CNV neuron. Neurons with 0 CNVs were excluded from this plot. (**F**) Percent of neurons containing CNVs plotted against the age of the individual from which they were gathered across all individuals (R^2^ = 0.5521; p = 0.00097). Lenient CNV detection thresholds were used for all of the above.

**Figure S3.**
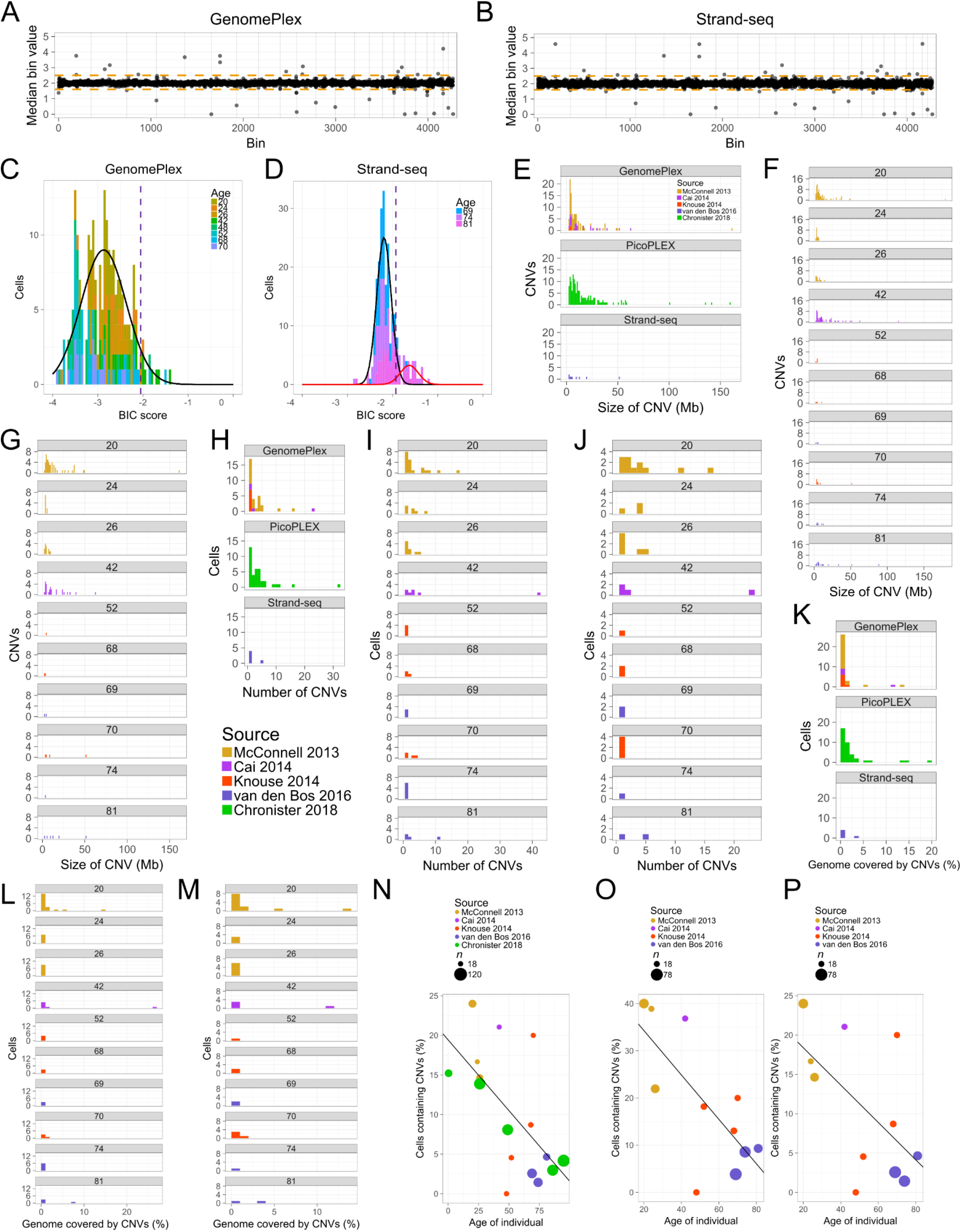
(related to Figure 3). Publicly available data shows similarities to PicoPLEX data under both CNV thresholds. (**A**) Median bin values for autosomal bins in GenomePlex data. Orange dashed lines represent bin exclusion thresholds of > 2.23 and < 1.80, as determined by Tukey’s outlier test. (**B**) Median bin values for autosomal bins in Strand-seq data. Orange dashed lines represent bin exclusion thresholds of > 2.32 and < 1.73, as determined by Tukey’s outlier test. (**C**) Histogram of BIC scores for GenomePlex datasets. Gaussian distribution fit to data is depicted in black. Dashed line indicates BIC cutoff of < −2.05. (**D**) Histogram of BIC scores for Strand-seq datasets. Gaussian distributions fit to data are depicted in black and red. Dashed line indicates BIC cutoff of < −1.93. (**E**) Histogram showing CNV sizes among neurons across each WGA method and data source. Stringent thresholds were used. (F and G) Histogram showing CNV sizes in each individual from previously published data. CNVs were detected using lenient thresholds (**F**) and stringent thresholds (G). (**H**) Histogram showing number of CNVs in CNV neurons across each WGA method and data source. Neurons with 0 CNVs were excluded from this plot. Stringent thresholds. (I and J) Histogram showing number of CNVs in CNV neurons in each individual from previously published data. Neurons with 0 CNVs were excluded from this plot. CNVs were detected using lenient thresholds (**I**) and stringent thresholds (J). (**K**) Histogram showing percent genome coverage by CNVs in CNV neurons across each WGA method and data source. Neurons with 0 CNVs were excluded from this plot. Stringent thresholds. (L and M) Histogram showing percent genome coverage by CNVs in CNV neurons in each individual from previously published data. Neurons with 0 CNVs were excluded from this plot. CNVs were detected using lenient thresholds (**L**) and stringent thresholds (M). (**N**) Percent of neurons containing CNVs plotted against the age of the individual from which they were collected (R^2^ = 0.3941; p = 0.0092). Stringent thresholds. (O and P) Percent of neurons containing CNVs plotted against the age of the individual using previously published data under lenient CNV thresholds (**O**) (R^2^ = 0.5315; p = 0.011) and stringent thresholds (**P**) (R^2^ = 0.3524; p = 0.054).

Published datasets performed WGA using GenomePlex (Cai et al., 2014; Knouse et al., 2014; McConnell et al., 2013), a DOP-PCR based approach, or Strand-seq (van den Bos et al., 2016), a “pre-amplification free” approach. We applied Tukey’s outlier test separately to GenomePlex and Strand-seq datasets to identify outlier bins for exclusion (Fig. S3A-B). This test resulted in the exclusion of 150 bins in male GenomePlex cells, 153 bins in female GenomePlex, 133 bins in male Strand-seq, and 135 bins in female Strand-seq. Across all sexes and WGA methods, including male PicoPLEX, 31 common outlier bins were identified, indicating a common source of noise/bias for these loci in the genome. Following outlier bin exclusion and segmentation, BIC cutoffs were calculated using the same approach as for the PicoPLEX data (Fig. S3C-D). 214 of 225 GenomePlex cells (95.1%) passed the cutoff of BIC < −2.05 while 191 of 233 Strand-seq cells (82.0%) passed the cutoff of BIC < −1.93 (Fig. 3A, B).

The general characteristics of CNVs were similar regardless of WGA approach or sex (Fig. 3C, D, E). The mean length of single cell CNVs was 14.8 Mb across all datasets, which is notably larger than most CNVs observed in bulk, germline human genomes (1–10kb, (MacDonald et al., 2014)). As with PicoPLEX data, CNV size was more variable among individuals than among datasets (Table 2). The average CNV size in GenomePlex individuals was 11.7 Mb, but ranged from 4.1 to 15.9 Mb among individuals. The average CNV size in Strand-seq individuals was 13.2 Mb but ranged from 4.3 to 17.7 Mb among individuals. Similar, albeit on average larger, CNV sizes were observed at the stringent threshold for all individuals (Fig. S3E). As observed in our PicoPLEX dataset (Fig. S1 L-M) the strongest predictor of CNV neuron prevalence in an individual was their age.

**Table 2.** Overview of reanalyzed publicly available data.

| Age | Sex | Cell type | Data source | WGA | BIC pass/total cells (%) | Cells with CNV(s) (%), lenient | Cells with CNV(s) (%), stringent | Total CNVs (lenient) | Total CNVs (stringent) |
| --- | --- | --- | --- | --- | --- | --- | --- | --- | --- |
| 20 | Female | Neuron | McConnell 2013 | GenomePlex | 50/50 (100) | 20 (40.0) | 12 (24.0) | 76 | 52 |
| 24 | Female | Neuron | McConnell 2013 | GenomePlex | 18/19 (94.7) | 7 (38.9) | 3 (16.7) | 20 | 9 |
| 26* | Male | Neuron | McConnell 2013 | GenomePlex | 41/41 (100) | 9 (22.0) | 6 (14.6) | 18 | 13 |
| 42 | Female | Neuron | Cai 2014 | GenomePlex | 19/26 (73.1) | 7 (36.8) | 4 (21.1) | 57 | 27 |
| 48 | Female | Neuron | Knouse 2014 | GenomePlex | 21/22 (95.5) | 0 (0) | 0 (0) | 0 | 0 |
| 52 | Male | Neuron | Knouse 2014 | GenomePlex | 22/22 (100) | 4 (18.2) | 1 (4.5) | 4 | 1 |
| 68 | Male | Neuron | Knouse 2014 | GenomePlex | 23/25 (92.0) | 3 (13.0) | 2 (8.7) | 4 | 2 |
| 69 | Male | Neuron | van den Bos 2016 | Strand-seq | 78/81 (96.3) | 3 (3.8) | 2 (2.6) | 3 | 2 |
| 70 | Male | Neuron | Knouse 2014 | GenomePlex | 20/20 (100) | 4 (20.0) | 4 (20.0) | 9 | 4 |
| 74 | Male | Neuron | van den Bos 2016 | Strand-seq | 70/80 (87.5) | 6 (8.6) | 1 (1.4) | 6 | 1 |
| 81 | Female | Neuron | van den Bos 2016 | Strand-seq | 43/72 (59.7) | 4 (9.3) | 2 (4.7) | 15 | 6 |

\*Same individual as 26 year old listed in Table 1.

^*^Same individual as 26 year old listed in Table 1.

The negative correlation between age and the percentage of CNV neurons remained significant upon addition of the publicly available data (Fig. 3F). Obvious confounding variables such as BIC score or post-mortem interval showed no similar correlation with age (Fig. S4). Plotting the entirety of the data, we found that the correlation became more significant by roughly an order of magnitude (R^2^ = 0.5521; p = 0.00097) (Fig. 3F). The correlation was also significant using the stringent threshold (R^2^ = 0.3941; p = 0.0092) (Fig. S3N). Although there were few young individuals in the published data set, we also tested whether the correlation was present in only the published datasets. With the lenient threshold, the correlation was significant (R^2^ = 0.5315; p = 0.011) (Fig. S3O); however, with the stringent threshold, the correlation was mildly insignificant (R^2^ = 0.3524; p = 0.054) (Fig. S3P).

**Figure S4.**
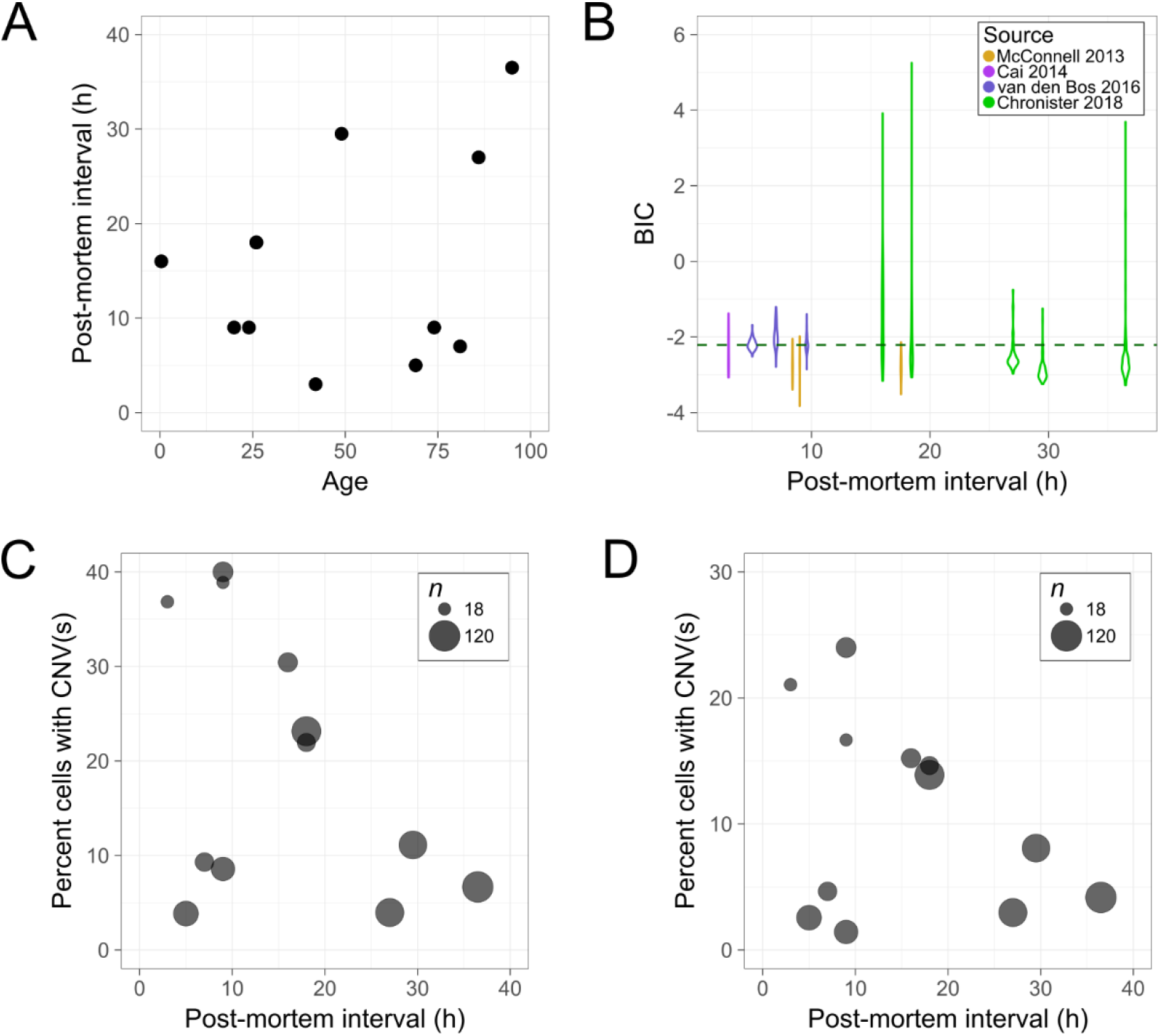
(Related to Figure 3). Post-mortem interval (PMI) does not impact sample quality or confound age-related trends. (**A**) Scatter plot of PMI and age information for 11 individuals studied in this paper. (**B**) Violin plots of BIC score and PMI for 11 individuals studied in this paper. The 26 year old individual studied in both McConnell 2013 and this paper has a PMI of 18 h and violins are shown for each group of cells analyzed. (C and D) Percent of neurons containing CNVs plotted against the PMI of the individual from which they were collected using lenient CNV thresholds (**C**) and stringent thresholds (D). The data points represent 11 of the individuals studied in this article, including the 26 year old for whom there are separate data points for the McConnell 2013 and Chronister 2018 datasets. The relationship was statistically insignificant in both cases.

Our observation that fewer CNV neurons are observed in aged individuals is in contrast to the concept of genosenium (Lodato et al., 2017), which states that the accumulation of somatic mutations over one’s lifetime is associated with aging-related cellular and molecular phenotypes. Thus, we also analyzed the number of CNVs per CNV neuron. Although most (62.8%) CNV neurons contained only 1 or 2 CNVs, the average number of CNVs per neuron in the dataset was 3.9 (Fig. 3D). Older CNV neurons, although rare, tended to have the most CNVs / neuron. Notably, the 20 year old individual had the most CNV neurons (40% CNV neurons, 3.8 CNVs/neuron), but the 49 year old (11.1% CNV neurons) had the most CNVs per neuron (10.3). Other individuals, such as the 24 year old and 86 year old, also support this trend; we detected 38.9% CNV neurons but 2.8 CNVs/neuron in the 24 year old individual and 4.0% CNV neurons but 5.0 CNVs/neuron in the 86 year old individual. Genosenium, if true, may operate on different somatic mutations in distinct ways.

### Long Gene Mosaicism and Neuronal Diversity

Brain CNVs, like all CNVs, occur because DNA repair is not perfect. Transcription and replication lead to DNA double-strand breaks (DSBs), which in turn sometimes lead to CNVs. Intriguingly, gene length increases susceptibility to transcriptional and replicative genomic stress. Genes encoded by more than 100kb of genomic sequence (i.e., long genes) tend to be neuronally expressed genes with roles in neuronal connectivity and synaptic plasticity (Zylka et al., 2015). Long genes also overlap with DNA fragile sites, and replicative stress can lead to large CNVs that encompass these loci (Wilson et al., 2015). Likewise, transcriptional stress leads to DNA DSBs in neurons (Madabhushi et al., 2015) and has a predominant effect on long gene transcript abundance (King et al., 2013). Recent studies link these observations to DNA DSBs during mouse neurodevelopment (Wei et al., 2016) and motivate the hypothesis that somatic mutations affecting long genes mediate the functional consequences of brain somatic mosaicism (Weissman and Gage, 2016).

The assembled atlas identified 129 CNV neurons and 8 CNV non-neurons that collectively harbor 522 neural CNVs (Tables 1 and 2). We performed overlap and random permutation analyses to determine if subsets of 93 candidate long genes (Fig. 4A) were associated with CNVs more frequently than expected by chance. Genomic regions (i.e., bins) that accumulated CNVs at an increased population-wide frequency compared to the rest of the genome were assessed at relevant thresholds (see methods), as were stringent and lenient thresholds, duplications, and deletions (Fig. 4B). Enrichment was assessed by generating 184 raw p values from each of 8 candidate gene lists was determined relative to 10,000 permutations of 23 different sets of genomic regions. After correcting for multiple hypothesis testing (Benjamini-Hochberg false discovery rate (FDR); 5% FDR cutoff), dataset-wide significance was observed for the entire candidate gene list, and, notably, with putative hot-spots that encompass 3 of the 4 common candidate genes, GPC6, NRXN3, RBFOX1 (Fig. 4C).

**Figure 4.**
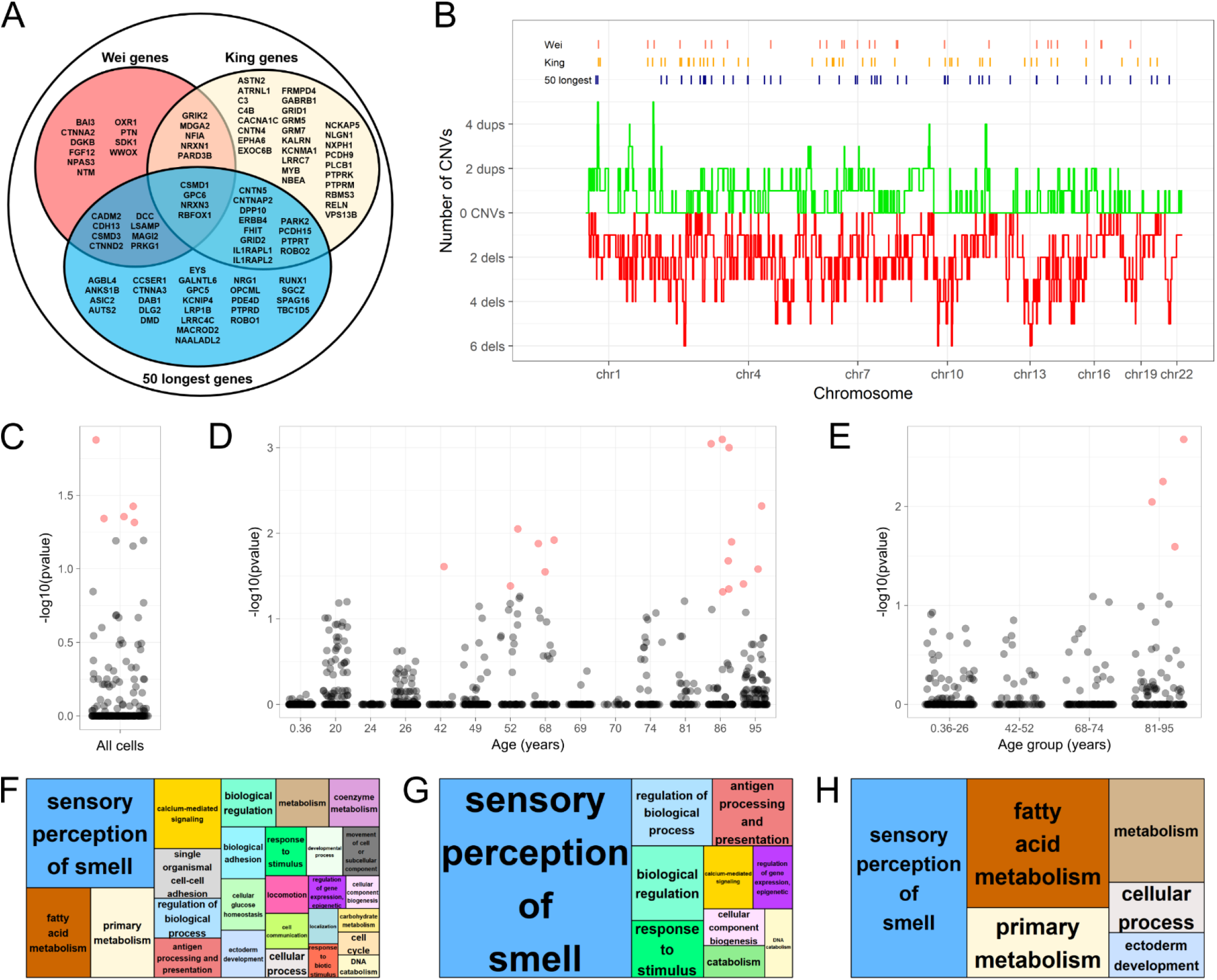
Long gene enrichment in mosaic CNVs. (**A**) Venn diagram of gene lists used for enrichment. (**B**) Summary of all CNVs detected in PicoPLEX and published data; red lines represent deletions while green lines represent duplications. Lenient CNV thresholds were used. (C-E) Enrichment results for hotspots from all cells (C), individuals (D), and age groups (E). P-values < 0.05 are colored in red. (F-H) Plots illustrating enriched categories of GO terms from REViGO analysis. The size of each box corresponds to the significance of the labeled GO category, and the largest boxes have the greatest overall significance. GO enrichment was found using PANTHER analysis of CNV-affected genes in all neural data (F), and age groups 68–74 (**G**) and 81–95 years old (H).

CNV neurons were rare in aged individuals so we assessed whether the genetic architecture of CNV neurons may also change during lifespan. We further tested each individual in the atlas for candidate gene enrichment using applicable thresholds for recurrent hot-spots (see methods) at stringent and lenient criteria (Fig. 4A). Significant corrected p-values were not observed in the youngest individuals, but were observed in aged individuals (Fig. 4D). When individuals were pooled based on age groups, significant p-values were observed only in the most aged group (Fig. 4E).

CNV-affected loci may not be restricted to long genes. We assembled a comprehensive list of all genes affected in the brain CNV atlas and used PANTHER (Mi et al., 2013) to calculate enrichment statistics and plotting scripts from REViGO (Supek et al., 2011) to visualize these data. Gene ontology categories associated with “sensory perception of smell” and “calcium-mediated signaling” were notably enriched in the atlas-wide gene set (Fig. 4F). When assessed by age group as in Fig. 4E, these enrichments were only detected in the aged groups (Fig. 4G, H).

## Discussion

Brain somatic mosaicism is a largely unexplored aspect of neuronal diversity. In the cerebral cortex, neuronal diversity is described in terms of electrophysiological properties (Contreras, 2004) and gene expression profiles (Lake et al., 2016) that are brought about by genetic programs (Lein et al., 2017). Current neurodevelopmental models implicitly assume that all somatic cells operate with identical genomes. In turn, population-based genetic studies of neurological disease typically sequence bulk blood DNA as a proxy for brain DNA. Somatic mutations affecting cell proliferation and survival pathways can alter neuronal diversity and lead to cortical overgrowth phenotypes ranging from hemimegalencephaly to focal dysplasia (Jamuar et al., 2014; Lee et al., 2012; Mirzaa et al., 2016; Poduri et al., 2012). Elevated levels of somatic mutations have also been associated with Rett Syndrome (Muotri et al., 2010), neurodegenerative disease (Bushman et al., 2015; Iourov et al., 2009; Lodato et al., 2017; McConnell et al., 2004), schizophrenia (Bundo et al., 2014), and altered behavior (Bedrosian et al., 2018). The Brain Somatic Mosaicism Network is an ongoing multi-site effort that aims to define how brain somatic mutations affect the genetic architecture of psychiatric disease (McConnell et al., 2017). However, the consequences of somatic mutations in neurotypical human brains remain a central unaddressed question. We report a comprehensive assessment of brain somatic CNVs to begin to discern how CNV neurons alter neuronal diversity in neurotypical human cerebral cortex.

We assembled an atlas of genomic data from 1285 single neural nuclei obtained from 15 neurotypical individuals. Given that 4 different laboratories have used 3 different WGA approaches to affirm the existence of CNV neurons, we developed objective read depth-based CNV detection sequentially. First, we identified optimal DNAcopy parameters that showed high sensitivity, high specificity, and minimal BIC scores on 6 test datasets, and applied a single WGA approach to >100 neurons from each of 5 neurotypical individuals. Second, we applied optimized DNAcopy to the entire 829 PicoPLEX WGA dataset to identify high quality libraries based on BIC scores and remove outlier genomic bins. Integer-like CNV segments were apparent in segment CN state distributions centered near CN = 1, 2, and 3. Third, we obtained and harmonized published datasets then employed optimized segmentation to single nucleus libraries generated using either GenomePlex or Strand-seq WGA approaches. Collectively, this approach yielded a final dataset comprised of 879 neuronal and 115 non-neuronal genomic libraries and identified Mb-scale CNVs in 129 neurons and in 8 non-neurons.

Salient features of the brain CNV atlas reveal that changes in the genetic architecture of neurotypical brain somatic CNVs are associated with an individual’s age. We found that CNV neuron prevalence was markedly diminished in aged individuals. This correlation was significant in both our new PicoPLEX dataset and the published GenomePlex/Strand-seq dataset. Based on the significant negative correlation (P = 0.00097) observed between CNV neuron frequency and age among these 15 individuals, we conclude that there is less than a 0.1% chance that the correlation observed simply reflects natural human variation, sampling idiosyncrasies, or unmeasured attributes of sample quality. By contrast, initial assessment of CNV location finds evidence for enrichment of a subset of long-genes and neurally-associated gene ontology categories only in aged brains. This may indicate that some somatic CNVs are more compatible with neural survival than others.

We provide the first evidence that a functional consequence of CNV neurons may be selective vulnerability to aging-related cell death. Age-related cognitive decline is associated with notable decreases in cerebral cortical thickness, myelination, and synapse number accompanied by *ex vacuo* enlargement in ventricular volume. Although neuronal cell death is generally considered to be minimal in the healthy mature brain, rates of ∼10% cerebral cortical neuron loss during adulthood are consistent with stereological counts in neurotypical individuals (Pakkenberg et al., 2003). The decline in CNV neuron prevalence that we observe between individuals <30 years old and individuals >70 years old is also strikingly consistent with ∼10% cortical neuron loss over a person’s adult lifetime. We conclude that the most parsimonious interpretation of these data is that many CNV neurons are selectively vulnerable to aging-associated atrophy. This finding highlights the unmet need for an increased understanding of human neuronal genome dynamics both during neurogenesis and among mature neurons. Human pluripotent stem cell-based models (Brennand et al., 2015) represent a straightforward means to this end.

## Acknowledgements

We thank J. Lannigan and M. Solga (UVa FACS core), Y. Bao (UVa Genome analysis and technology core), and A. Koeppel (UVa Bioinformatics core) for contributed expertise. We thank P. Lansdorp (UBC) for prompt sharing of unpublished metadata, and F.H. Gage (Salk) and J.V. Moran (UM) for critical feedback. Human tissue was obtained from the National Institute for Child Health and Human Development (NIH) Brain and Tissue Bank for Developmental Disorders at the University of Maryland, Baltimore, MD, contract HHSN2752009000011C, ref. no. N01-HD-9-011. Data are available through the Brain Somatic Mosaicism Portal (synapse.org/BSMN) and stored at the NIH data archive (Accession xxxxx). Funding to MJM (U01 MH106882), to DJW (U01 MH106893), and to MJM and SB (U01 MH106882-03S1) supported this work. IEB received support from the McDonnell foundation, and WDC received support from T32 GM008136-30.

## Materials And Methods

### Single nucleus isolation and WGA

Single nucleus isolation was performed as before (McConnell et al., 2013) and as described in detail (Wierman et al., 2017). Post-mortem cortex was stored at −80C and fragmented on dry ice in a pre-chilled pestle. Fragments (∼0.3 mg) were collected in lysis buffer and homogenized. We purified nuclei on an iodyxonol cushion by ultracentrifugation. We immunostained nuclei overnight (4 °C) with 1:250 diluted mouse monoclonal anti-human NeuN IgG Alexa-Fluor 555 Conjugate clone A60 (EMD Catalog #MAB377A5; Millipore, Burlington, MA). We sorted nuclei into 8-well PCR tube strips and applied PicoPlex WGA according to manufacturer’s instructions (Rubicon Genomics, Ann Arbor, MI). We identified successful WGA by gel electrophoresis on 1% agarose gel and then purified each positive reaction with QIAquick PCR purification columns (Qiagen, Germantown, MD). DNA was quantified using HS DNA assay (ThermoFisher, Waltham, MA). We size-selected pooled libraries 450 bp to 800 bp DNA fragments in 7.5% acrylamide and then electro-eluted DNA into 1% agarose prior to purification using QIAquick gel extraction kits (Qiagen). We quantified the DNA concentration of all libraries by HS DNA Qubit 2.0 (ThermoFisher) and diluted to 6 nM prior to sequencing on an Illumina (San Diego, CA) platform.

### Analysis of single cell sequencing data

Sequence reads from Illumina were trimmed of PicoPLEX primers using the fastx_trimmer command (hannonlab.cshl.edu/fastx_toolkit/). Reads were then aligned to the human genome (version hg19) with BWA-aln V0.7.12 using default options (Li and Durbin, 2009). Duplicates were removed using MarkDuplicates (Picard tools V1.105, broadinstitute.github.io/picard). Using a 40-mer mappability track (UCSC, wgEncodeCrgMapabilityAlign40mer.bigWig) to determine uniquely mappable bases, we divided the genome into 4,505 dynamically sized genomic bins, each containing 500kb of mappable sequence. The mean bin size was 687 kb. Read counts for each bin were determined by Bedtools V2.17.0 coverageBed (Quinlan and Hall, 2010). To avoid read count bias arising from GC content, bins were grouped into 16 roughly equal size groups according to GC percentage and each read count within a GC group was divided by the median read count of the group and multiplied by two.

Following analysis of several hundred single cell datasets, we observed that certain genomic bins were consistently above or below the euploid state, most likely due to biases arising from alignment or artifacts generated during WGA. To avoid biases in segmentation introduced by these outlier bins, namely false positive and false negative CNVs, we used Tukey’s Outlier Test on the median log-copy number values of all 4,505 bins in a sex and WGA-specific fashion, resulting in between 101 and 153 bins to be excluded from segmentation, depending on the sex of the individual and WGA used. In addition to the bins excluded by the outlier test, two Y chromosome bins were manually excluded from female Strand-seq dataset. Single cell datasets were segmented using DNAcopy (Seshan and Olshen, 2017), an R package (www.R-project.org) that implements circular binary segmentation (CBS) to detect copy number “changepoints” in genomic data. DNAcopy was run on the normalized bin data using parameters alpha=0.001, undoSD=0, and min.width=5.

### BIC scoring and filtering

To determine which samples were of sufficient quality to merit further analysis, we implemented Bayesian Information Criterion (BIC)(Schwarz, 1978)

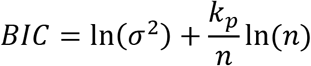

as a scoring metric where σ^2^ is the variance of the data points about their respective segment mean, kp is the number of segments, and n is the number of bins (4,505). BIC rewards a segmentation that avoids under-fitting the data (*i.e.*, allowing high variance of data within segments) and over-fitting the data *(i.e.*, creating too many segments); thus, a cell with properly fitted data will have a low BIC score as a result.

To define threshold BIC scores for inclusion in further analysis, a histogram of BIC scores was generated for each WGA method. Using the R package mixtools (Benaglia et al., 2009), we fit two Gaussian distributions to the PicoPLEX and Strand-seq histograms, and one Gaussian distribution to the GenomePlex histogram due to its displaying a single mode. Using the Gaussian distribution with the lower mean (or, in the case of GenomePlex, the lone distribution), we set the threshold BIC score for inclusion to correspond to p=0.05 on the upper tail. For PicoPLEX, cells scoring below −2.21 were selected for further analysis (Fig. S1G); for GenomePlex, the threshold was −2.05 (Fig. S3C); for Strand-seq, the cutoff was −1.93 (Fig. S3D).

### Defining CNVs

Because DNAcopy does not assign integer copy number values to the segments it outputs, it was necessary to define threshold copy number values for a segment to be considered a CNV. We set CNV thresholds by evaluating autosomal segments of sizes ranging from 5, our minimum number of bins required to call a CNV, and 45 bins, a length smaller than the shortest autosome, resulting in a distribution of copy numbers excluding the high number of whole chromosomes at or near copy number two. The segments were plotted by copy number value in a histogram, and we fit a three-Gaussian mixture model using mixtools (Benaglia et al., 2009) and plotted the resulting mixed Gaussian model of three distributions centered at the local peaks near copy number 1, 2, and 3. Using the central Gaussian, centered near 2, we calculated two sets of thresholds: the stringent thresholds, 2.80 and 1.14, the result of a cumulative two-tailed probability of 0.01; and the lenient thresholds, 2.60 and 1.34, determined by a two-tailed cumulative probability of 0.05. Throughout our CNV analyses, we used both thresholds in order to give a wider range of possible results and avoid any influence of false positive or false negative CNVs. Additionally, CNVs were required to be at least 5 bins in length, and X and Y were not examined for CNVs. Following the calculation of BIC and CNV thresholds, it was discovered that 2 cells in our analysis of 829 PicoPLEX cells contained female (XX) copy number profiles, and thus were excluded from downstream analysis, including CNV detection, gene set enrichment, and Gene Ontology (GO) term analysis.

### Test data simulation

DNAcopy is prone to calling CNV events using correlated noise as input (Muggeo and Adelfio, 2011), so we sought to determine the degree to which our single cell data contains correlated noise. We selected six single cells to simulate test data. In our “NULL model”, we assumed that the data for our six cells were described by correlated noise and Gaussian noise about the euploid copy number state, and contained no real CNV events. In the “alternative model”, we assumed that the six cells were described by real CNV events identified by DNAcopy and residual correlated and Gaussian noise. We simulated 200 test data cells for each of the cells under each of the models and found that the CNVs produced by the NULL model rarely matched those of the original cell in size or divergence from 2, leading us to conclude that the ALT model was a more accurate representation of our data. To determine the best DNAcopy segmentation parameters, we used simulation data of the “alternative model” to explore values of alpha ranging from 0.05 to 10^−5^, undoSD ranging from 0 to 5, and min.width ranging from 2 to 5, and calculated a BIC score for each. We also tested the performance of each segmentation using a receiver operating characteristic (ROC) curve to determine the parameters at which sensitivity was maximized and false positive CNVs were minimized (Fig. S1E). We concluded that the best parameters for segmentation were alpha=0.001, undoSD=0, and min.width=5.

### Gene set enrichment

To explore the biological significance of large scale CNVs in the brain, we sought to follow up on the work of two papers that identified lists of long genes that may be predisposed to DNA breaks in mice (King et al., 2013; Wei et al., 2016). Building upon this idea, we also examined a third list containing the 50 longest human genes obtained using biomaRt (Smedley et al., 2015). These gene lists shared genes in common with one another, and all told, there were ultimately seven gene lists drawn from one, two, or three of the original lists, as well as an eighth gene list containing all 93 genes gathered from all three sources. To test for enrichment, we collected hotspot coordinates corresponding to a minimum number of CNV events, which varied depending on the subset of data being tested and the CNV threshold used. For each data subset, we set hotspot cutoffs for the required number of CNVs such that no cutoff should lead to greater than 20% coverage of the human genome. The resulting cutoffs for each subset using lenient CNV thresholds were as follows: in the full combined dataset, hotspots were defined as regions containing 5–8 CNVs, 6–8 CNVs, 7–8 CNVs, 8 CNVs, 4–6 deletions, 5–6 deletions, 6 deletions, 2–5 duplications, 3–5 duplications, 4–5 duplications, and 5 duplications; we also defined two “coldspot” sets of regions based on coordinates which were affected by either 0 CNVs or 0 deletions. In the 0.36 year old individual, hotspots were defined as regions with 1–3, 2–3, and 3 CNVs, 1–2 and 2 deletions, and 1–2 and 2 duplications; in the 20 year old, 2–4, 3–4, and 4 CNVs, 2 deletions, and 1–2 and 2 duplications; in the 24 year old, 1 CNV, 1 deletion, and 1 duplication; in the 26 year old NeuN+ cells, 2–5, 3–5, 4–5, and 5 cNvs, 1–3, 2–3, and 3 deletions, and 1–4, 2–4, 3–4, and 4 duplications; in the 26 year old NeuN- cells, 1 duplication; in the 42 year old, 2 CNVs, 2 deletions, and 1 duplication; in the 49 year old NeuN+ cells, 2–3 and 3 CNVs, 2–3 and 3 deletions, and 1 duplication; in the 49 year old NeuN- cells, 1 CNVs, 1 deletion, and 1 duplication; in the 52 year old, 1–2 and 2 CNVs, 1–2 and 2 deletions, and 1 duplication; in the 68 year old, 1 CNV, 1 deletion, and 1 duplication; in the 69 year old, 1 CNV, 1 deletion, and 1 duplication; in the 70 year old, 1 deletion; in the 74 year old, 1–2 and 2 CNVs, 1–2 and 2 deletions, and 1 duplication; in the 81 year old, 1 CNV, 1 deletion, and 1 duplication; in the 86 year old NeuN+ cells, 2 CNVs, 1 deletion, and 1 duplication; in the 86 year old NeuN- cells, 1 CNV, 1 deletion, and 1 duplication; in the 95 year old, 1–3, 2–3, and 3 CNVs, 1–2 and 2 deletions, and 1 duplication. In the age group of individuals aged 0.36 to 26 years, hotspots were defined as regions containing 3–5, 4–5, and 5 CNVs, 2–5, 3–5, 4–5, and 5 deletions, and 2–4, 3–4, and 4 duplications; in the age group of individuals aged 42 to 52, 3–4 and 4 CNVs, 3–4 and 4 deletions, and 1 duplication; in the age group of individuals aged 68–74, 1–2 and 2 CNVs, 1–2 and 2 deletions, and 1–2 and 2 duplications; in the age group of individuals aged 81–95, 2–4, 3–4 and 4 CNVs, 2–3 and 3 deletions, and 1–2 and 2 duplications.

Under stringent thresholds, the hotspots tested were as follows: in the full combined dataset, hotspots were defined as regions containing 4–6 CNVs, 5–6 CNVs, 6 CNVs, 3–5 deletions, 4–5 deletions, 5 deletions, 2–4 duplications, 3–4 duplications, and 4 duplications; we also defined one “coldspot” set of regions based on coordinates which were affected by 0 CNVs. In the 0.36 year old individual, hotspots were defined as regions with 1–3, 2–3, and 3 CNVs, 1 deletion, and 1–2 and 2 duplications; in the 20 year old, 2–4, 3–4, and 4 CNVs, 1–2 and 2 deletions, and 1–2 and 2 duplications; in the 24 year old, 1 deletion; in the 26 year old NeuN+ cells, 1–3, 2–3, and 3 CNVs, 1–2 and 2 deletions, and 1–2 and 2 duplications; in the 42 year old, 1 deletion; in the 49 year old NeuN+ cells, 2 deletions; in the 49 year old NeuN- cells, 1 deletion; in the 52 year old, 1 deletion; in the 68 year old, 1 CNV, 1 deletion, and 1 duplication; in the 69 year old, 1 cNv, 1 deletion, and 1 duplication; in the 70 year old, 1 deletion; in the 74 year old, 1 duplication; in the 81 year old, 1 CNV, 1 deletion, and 1 duplication; in the 86 year old NeuN+ cells, 1 deletion and 1 duplication; in the 86 year old NeuN- cells, 1 deletion; in the 95 year old, 1–3, 2–3, and 3 CNVs, 1–2 and 2 deletions, and 1 duplication. In the age group of individuals aged 0.36 to 26 years, hotspots were defined as regions containing 2–5, 3–5, 4–5, and 5 CNVs, 2 deletions, and 1–3, 2–3, and 3 duplications; in the age group of individuals aged 42 to 52, 2–3 and 3 deletions; in the age group of individuals aged 68–74, 1–2 and 2 CNVs, 1 deletion, and 1–2 and 2 duplications; in the age group of individuals aged 81–95, 2–4, 3–4 and 4 CNVs, 2–3 and 3 deletions, and 1 duplications.

For each set of hotspots, we randomly shuffled the hotspot loci within the genome 10,000 times to generate a NULL model of CNV coverage. We then determined the number of genes of interest found in the actual hotspots and the range of genes of interest found in the NULL model to generate enrichment p-values, which were then FDR-corrected using the Benjamini-Hochberg method. Owing to the considerable inter-dependence among hotspot sets being tested, these p-values were FDR-corrected within stratified sub-groups; for example, p-values for hotspots derived from individuals were corrected separately from p-values for hotspots derived from age groups. Likewise, p-values for hotspots defined as regions of 3–5 CNVs were corrected separately from regions defined as regions of 2–5, 4–5, or 5 CNVs.

### Gene Ontology (GO) term analysis

We compiled a list of genes with genomic coordinates overlapping each CNV and separated these lists by individual and, where applicable, cell type. These lists of CNV-affected genes were submitted to PANTHER (pantherdb.org) to determine if any GO terms were enriched. The resulting GO terms and corresponding p-values were then submitted to REViGO (revigo.irb.hr) to aid visualization via downloadable plotting scripts.

